# The Photoswitchable Cannabinoid *Azo*-HU308 Enables Optical Control of Ca^2+^ Signaling in Pancreatic β-Cells via a Non-CB2 TRPC Channel Mechanism

**DOI:** 10.1101/2025.03.21.644674

**Authors:** Alexander E.G. Viray, James A. Frank

## Abstract

**Background and Purpose:** Ca^2+^ plays a critical role in regulating insulin secretion from pancreatic β-cells, a process modulated by various cell surface receptors, including cannabinoid receptors (CBRs). However, our understanding of cannabinoid signaling in β-cells is complicated by the intricate pharmacology of cannabinoid ligands and their inherent hydrophobicity, which hinders precise control of receptor activation. This study aims to investigate the effects of the light-activatable CB2 receptor agonist, *azo*-HU308, on β-cell Ca^2+^ dynamics.

**Experimental Approach:** We employ fluorescent Ca^2+^ imaging in INS-1 832/13 (INS-1) β-cells to measure Ca^2+^ transients induced by *azo*-HU308 and photoactivation with UV-A light. We then apply a pharmacological screen using a various CBR and TRP channel antagonists to determine the mechanism by which *azo*-HU308 enables optical control of β-cell Ca^2+^ levels.

**Key Results:** We observed that *azo-*HU308 triggers a robust increase in intracellular Ca^2+^ when isomerized to the *cis*-form with UV-A light. The effect was repeatable over multiple cycles of irradiation and gradually desensitized on each sequential UV-light pulse. A pharmacological screen determined that the Ca^2+^ increase was not due to CB2 receptor activation and Ca^2+^ release from intracellular stores, but rather influx of extracellular Ca^2+^ through TRPC channels.

**Conclusions and Implications:** *azo-*HU308 enables robust, repeatable stimulation of Ca^2+^ in INS-1 pancreatic β-cells when triggered by UV-A light. This study presents a novel tool to optically control β-cell Ca^2+^ dynamics, and sheds light on a new mechanism by which synthetic cannabinoids affect Ca^2+^ signaling through non-GPCR targets.

## INTRODUCTION

Ca^2+^ regulates intracellular signaling and the secretion of hormones and neurotransmitters from excitable cells. In pancreatic β-cells, Ca^2+^ controls the release of insulin, a key hormone that regulates glucose homeostasis^1^. After a meal, blood glucose is elevated and is transported into the β-cell where it is metabolized to ATP. This closes ATP-sensitive K^+^ channels (K_ATP_) and depolarizes the β-cell, triggering Ca^2+^ influx through voltage-dependent Ca^2+^ channels and subsequent insulin release. In addition, β-cell Ca^2+^ is regulated by myriad receptor pathways, including ion channels and G protein-coupled receptors (GPCRs). These Ca^2+^ fluctuations are oscillatory in nature, and their intensity and frequency correlates insulin release^2^. Dysfunction in β-cell Ca^2+^ signaling and GSIS leads to type 2 diabetes mellitus (T2DM), a disease whose prevalence is rising throughout the world at epidemic rates. As such, novel approaches to modulate β-cell Ca^2+^ levels and insulin release are urgently needed to address this epidemic.

Small molecule therapeutics that treat diabetes by promoting insulin release include sulfonylureas, which block K_ATP_ channels^3–6^, and approaches that modulate the β-cell’s intracellular Ca^2+^ concentration ([Ca^2+^]_i_) through alternative receptor pathways, including GPCR targets, have gained significant therapeutic interest^7^. For example, the GPR40 agonist TAK-875 improved glycemic control in patients, but was pulled from clinical trials due to toxicity concerns^8,9^. More recently, glucagon-like-peptide-1-receptor (GLP1R) agonists have demonstrated an improved safety for treating metabolic disease, and are increasingly being prescribed for T2DM^10^. Importantly, β-cells express nearly 300 GPCRs that modulate [Ca^2+^]_i_^11,12^, and these represent unexplored therapeutic targets that could be eventually be leveraged as new therapeutic targets for T2DM.

One such GPCR family are the cannabinoid receptors (CBRs), which are activated by endocannabinoid lipids and phytocannabinoids found in marijuana. Cannabinoid CB1 and CB2 receptors (CB1R and CB2R) are expressed in the human pancreas and effect β-cell excitability^13,14^. Additionally, non-canonical “orphan” CBRs—like GPR55^15,16^—affect insulin release from β-cells *in vitro* and in rodent models^17,18^. Unfortunately, cannabinoid-based therapeutics have had difficulty translating to the clinic, likely because we have limited understanding of how CBRs affect β-cell function. This is due to the hydrophobic nature and of cannabinoid ligands; these are insoluble in physiological solutions, which makes their application to cells slow and irreversible^19,20^. Cannabinoids also have complex pharmacology, and act through multiple GPCR and ion channel targets^21–23^. To deconvolute the precise roles of CBR signaling in modulating β-cell excitability and leverage this important signaling system for therapeutic development, we require tools that enable greater spatiotemporal resolution over cannabinoid signaling pathways.

Photopharmacology is an emerging approach to address these challenges by enabling optical control of biological processes with light-sensitive small molecules, including photocaged and photoswitchable probes^24^. Photocaged ligands have their activity blocked by a photo-labile protecting group; where a flash of light irreversibly releases the active compound. Photocages for Ca^2+^ and other ligands, including endocannabinoids^16,25,26^, have been applied to optically control β-cell [Ca^2+^]_i_^27,28^. Alternatively, photoswitchable probes are ligands that contain a photo-isomerizable motif, most commonly an azobenzene, that switches between the *trans*- and *cis*-confirmations using two different wavelengths of light ^29,30^. The change in molecular structure through cis/trans isomerization tunes the potency and efficacy of the ligand, enabling reversible, optical control of receptor activation with superior kinetics compared to standard pharmacology. Numerous photoswitches—including sulfonylureas or endogenous lipids—have been shown to enable dynamic manipulation of [Ca^2+^]_i_ in cultured β-cells and pancreatic islets^31–36^. Although our group and others have developed an array of photoswitchable cannabinoids to modulate both CB1R^37–39^ and CB2R ^40,41^; to date, they have not been applied to β-cells.

Recently, our group developed a photoswitchable ligand based on the CB2R-selective synthetic cannabinoid, HU308^42^. We incorporated an azobenzene into HU308’s alkyl chain to afford a photoswitchable ligand, *azo*-HU308-3 (now coined *azo*-HU308) (**Fig. 1A**). Using a Ca^2+^ imaging assay in AtT-20 cells overexpressing CB2R [AtT-20(CB2) cells], we demonstrated that *azo*-HU308 enables reversible, optical control of [Ca^2+^]_i_ when photoswitched to *cis*, and that the mechanism proceeded by CB2R and phospholipase C, and subsequent Ca^2+^ release from intracellular stores. As a follow-up to this study; here, we present the application of *azo-*HU308 to optically control [Ca^2+^]_i_ in cultured pancreatic β-cells. We show that *azo*-HU308 robustly elevates intracellular Ca^2+^ in β-cells when isomerized to *cis* with UV-A light and permits multiple rounds of stimulation. Using a pharmacological screen to determine mechanism-of-action, we determined that the Ca^2+^ influx was not CB2R-mediated, but rather involves TRPC ion channels and Ca^2+^ influx across the plasma membrane. This study presents a novel tool to modulate β-cell Ca^2+^ dynamics using light, and sheds light on a new mechanism by which synthetic cannabinoids may modulate Ca^2+^ influx through non-GPCR targets.

**FIGURE 1.**
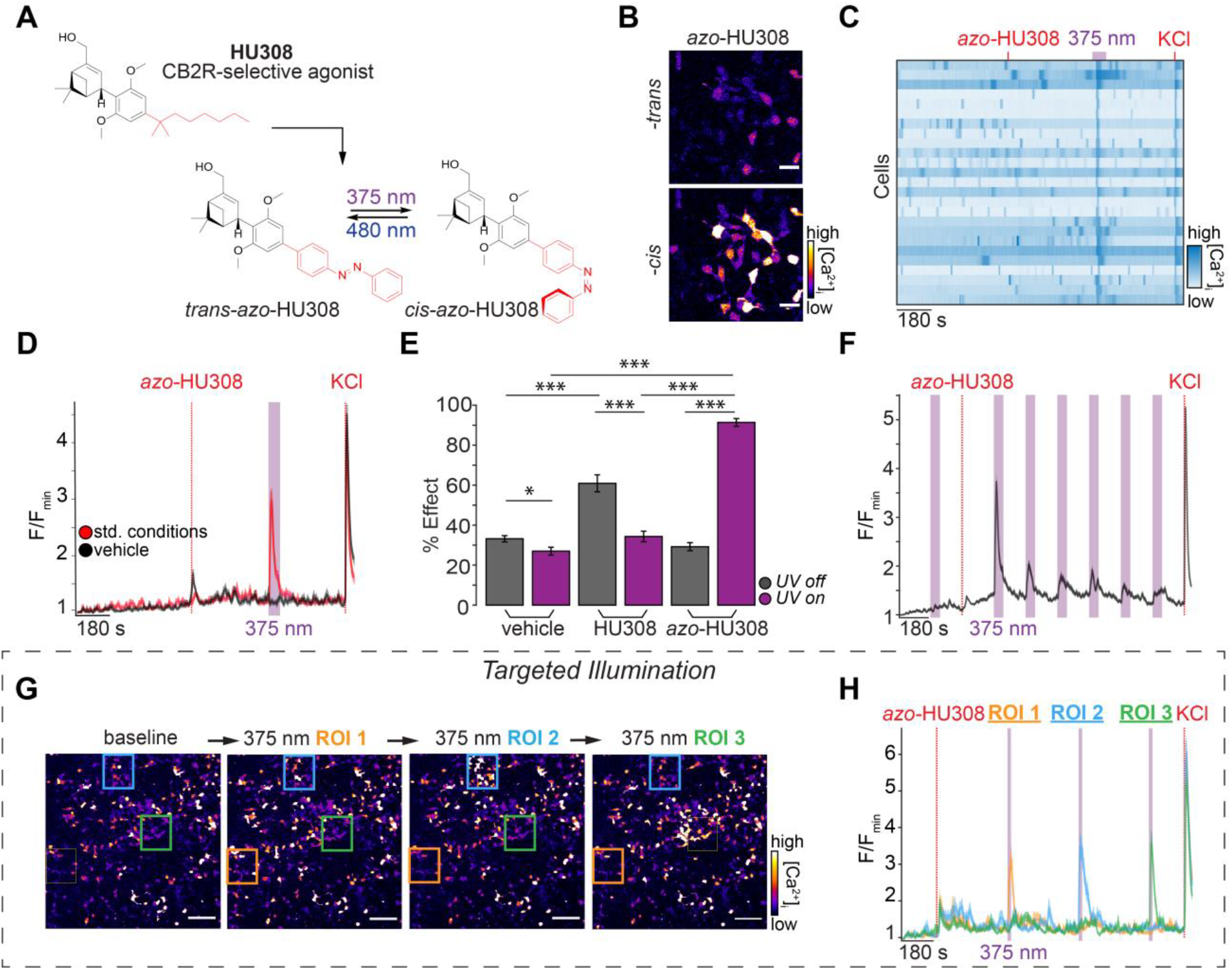
*Azo*-HU308 enables optical control of Ca^2+^ in INS-1 β-cells. (**A**) Chemical structure of HU308, which inspired design for the photoswitch *azo*-HU308. *azo*-HU308 isomerizes between *trans* and *cis* under blue and UV-A light, respectively. (**B-H**) INS-1 cells were transfected with R-GECO and imaged by confocal microscopy, then treated with *azo*-HU308 (20 μM) and 375 nm light pulses. Displayed are: (**B**) Representative images of R-GECO fluorescence in cells treated with *azo*-HU308 before (top) and after (bottom) 375 nm irradiation, which increases Ca^2+^ levels. (**C**) Heat map showing Ca^2+^ traces from 25 representative cells treated with *azo*-HU308 and irradiated with 375 nm light. (**D**) Averaged Ca^2+^ traces showing that *azo*-HU308 robustly stimulated Ca^2+^ during UV-A irradiation (red, N = 170, T = 4) and that vehicle (0.1% DMSO) had no effect (black, N = 203, T = 4). (**E**) Summary bar graph showing the effect on Ca^2+^ during compound addition (gray bars) and 375 nm irradiation (purple bars); comparing vehicle (N = 203, T = 4), HU308 (20 μM, N = 150, T = 3), and *azo*-HU308 (N = 170, T = 4). (**F**) Averaged R-GECO fluorescence plot demonstrating *azo*-HU308’s effect decays over multiple irradiations (N = 300, T = 3). (**G**) Representative images showing R-GECO fluorescence and three regions of interest (ROIs) targeted with 375 nm light during the Ca^2+^ imaging assay in H (N = 10, T = 3). (**H**) Averaged Ca^2+^ trace from cells in each of the ROIs in G, showing that *azo*-HU308 robustly stimulate Ca^2+^ release in only the cells that were irradiated (N = 10 per ROI, T = 3). Error bars = mean ± s.e.m. *P < 0.05, **P < 0.01, ***P < 0.001, ns = not significant.

## METHODS

### Compounds

*Azo*-HU308 (*azo*-HU308-3) was synthesized as previously described^43^. HU308 (Cayman Chemical #90086), AM630 (Cayman Chemical, #1006974), SR 144528 (Cayman Chemical, #9000491) JWH 133 (Cayman Chemical, #10005428), Rimonabant hydrochloride (Sigma-Aldrich, #SML0800), CID16020046 (Tocris, #4959), Xestospongin C (Cayman Chemical, #64950), U73122 (Tocris, #1268), YM254890 (Tocris, #7352), Pertussis Toxin (Cayman Chemical, #19546), NF449 (Cayman Chemical, #13324), 2-APB (Tocris, #1224), Capsazepine (Cayman Chemical, #10007518), SAR7334 hydrochloride (Cayman Chemical, #28292), SKF 96365 hydrochloride (Cayman Chemical, #10009312), ML-204 (Cayman Chemical, #15626), YM-58483 (Cayman Chemical, #13246) and Pyr 10 (Tocris, #6941) were obtained from commercial suppliers. Compounds were dissolved as stock solutions in DMSO, aliquoted and stored at-20 °C until use.

### Cell Culture Media and Solutions

**INS-1 media** contains: RPMI 1640 with L-glutamine (Gibco, #11875-093) with 10% FBS, 1:100 Penicillin-strep (5000 units/mL, Gibco, #15070) and 10 mM HEPES (Fisher, #BP310-500), 1 mM sodium pyruvate (Alfa Aesar, #A11148). INS-1 media was filtered and distributed into 50 ml aliquots. 50 μM 2-mercaptoethanol diluted in H_2_O (BME, Sigma, #M3148) was added fresh to each 50 ml aliquot prior to use.

**INS-1 imaging buffer** contains (in mM): 185 NaCl, 1.2 CaCl_2_, 1.2 MgCl_2_, 1.2 K_2_HPO_4_, 20 HEPES. Adjusted to pH 7.4 with NaOH. D-glucose was supplemented at 20 mM.

**Phosphate buffered saline (PBS)** contains (in mM): 320 Na_2_HPO_4_ (Fisher, #BP332-500), 80 Na(PO_4_H_2_)•H_2_O (Fisher, #S369-1). Adjusted pH to 7.4 with NaOH.

### Cell Culture

INS-1 832/13 cells^44^ were grown in INS-1 media and incubated at 37 °C and 5% CO_2_. Cells were used between passages 65-80. For live-cell Ca^2+^ imaging, INS-1 cells were plated at a density of 100,000 cells/well on 8-well glass bottom chambered coverslips (Ibidi, #0827-90 or Cellvis, #C8-1.5H-N). 18-24 h later, the cells were starved in Opti-MEM™ (250 μl) for 2 h before adding the transfection mixture containing (per well): 50 μl Opti-MEM™, 1 μl Lipofectamine-2000 (Fisher Scientific, #11668019) and 250 ng R-GECO cDNA. The cells were then incubated at 37 °C and 5% CO_2_ for 18-24 h before exchanging the transfection mixture with INS-1 media. Microscopy experiments were performed 60-72 h post-transfection.

### Confocal Microscopy

Live cell imaging was performed on an Olympus Fluoview 1200 laser scanning confocal microscope at 37 °C and 5% CO_2_. Videos were acquired with a 20×/0.75 NA objective (Olympus UPlanSApo) at 512×512-pixel resolution and scan rate of 4 s/frame. R-GECO excitation was performed with a 559 nm laser at low laser power (1-3%) and emission was collected at 570-670 nm. Compounds stock solutions (1000× in DMSO) were incubated for 10 s with 10% Pluronic F-127 (Tocris #6253) in DMSO (1:1 v:v ratio with compound solution). The mixture was then diluted 1000× with imaging buffer and pipetted directly into the imaging well. Vehicle controls were performed using an identical procedure, adding a final concentration of DMSO (0.2 vol%) and Pluronic F-127 (0.01 vol%). For co-application experiments with receptor antagonists, the drugs were applied to the imaging buffer and fluorescent images were acquired immediately upon thermal equilibration of the plate in the environment chamber, typically ~10 min after application.

Photoswitching was performed with a 375 nm laser (PicoQuant, PDL 800-D, ~95 μW output from the objective) triggered using the quench function in the Olympus software. KCl (30 mM) was added at the end of each experiment to elicit maximum R-GECO fluorescence intensity for normalization. Fluorescence intensity values in each experimental video were collected and analyzed with ImageJ^45^ and the data was processed and plotted in Excel and MATLAB using in-house written scripts (available on request). Oval-shaped regions were manually drawn over each cell to measure the fluorescence in ImageJ.

### Data Analysis and Statistical Methods

All data is presented as mean±s.e.m. For Ca^2+^ imaging experiments, the figure captions report the total number of individual cells (N), and the number of independent trials (T, biological replicates). Statistical significance was assessed using the Data Analysis package in MATLAB (Mathworks). For comparison between two groups, a Student’s paired t-test (two-sided) was used, with significance threshold placed at *P<0.05, **P<0.01, ***P<0.001, ns = not significant.

For bar graph analysis of R-GECO intensity (e.g. Fig. 1E), the photoswitching “Percent effect (%)” values were calculated as change in fluorescence intensity within 60 s after addition of the *trans* isomer; or for *cis-*, the maximum intensity during irradiation. These values are normalized to the range calculated between minimum baseline fluorescence intensity (first 50 frames) and the maximum intensity generated after KCl addition. These values were calculated for each cell, then calculated as a mean across all cells and plotted as a bar graph for comparison between conditions.

## RESULTS

### *Azo*-HU308 Enables Reversible Optical Control of Ca^2+^ influx in INS-1 pancreatic β-cells

Motivated by studies that showed that CBR signaling modulates Ca^2+^ in β-cells^25,26,46^ and our finding that *azo*-HU308 reversibly controls Ca^2+^ levels via CB2R and Ca^2+^ release from intracellular stores^43^, we asked whether *azo*-HU308 could optically modulate [Ca^2+^]_i_ in cultured β-cells. We chose the rat insulinoma INS-1 832/13 (INS-1) cells because they exhibit glucose-stimulated Ca^2+^ oscillations that couple to insulin release^44^, and little is known about CBR signaling in INS-1 cells. INS-1 cells were transfected with the genetically-encoded Ca^2+^ sensor R-GECO^47^, which allowed us to monitor [Ca^2+^]_i_ levels in real-time by fluorescence microscopy (**Fig. 1B, top**). We observed robust Ca^2+^ oscillations when the cells were exposed to elevated glucose (20 mM) solution (**Fig.1C**). Application of *trans*-*azo*-HU308 did not affect [Ca^2+^]_i_ (**Fig.1C, and Fig. 1D, red**); but strikingly, when we irradiated the cells with 375 nm UV-A light (~95 μW laser irradiation) to isomerize *azo*-HU308 to *cis* we observed a robust [Ca^2+^]_i_ burst across the cells (**Fig. 1B bottom, Fig. 1C, Fig. 1D red, Fig. 1E**). The Ca^2+^ increase was intense but short-lived and decayed rapidly before the photostimulation period was finished. The effect of *azo*-HU308 was dose-sensitive, and the [Ca^2+^]_i_-stimulating effect continued to increase up to 20 μM (**Fig. S1B and Fig. S1C**). In control experiments, the parent molecule HU308^42^ caused a similar Ca^2+^ response immediately upon addition, but was not effected by 375 nm irradiation (**Fig. S2**). In vehicle control experiments, where DMSO (0.1%) was added in the absence of *azo*-HU308, there was no change to [Ca^2+^]_i_ upon addition or irradiation (**Fig. 1D, black**). This confirms that the effect was not directly caused by the irradiation or an artefact of compound addition.

To test the reversibility of *azo*-HU308’s effects, 375 nm photostimulation was repeated over six cycles. We observed a Ca2+ increase on each irradiation, however the magnitude decayed on each subsequent stimulation, which indicates that receptor desensitization was likely occurring (**Fig. 1F**). To highlight the outstanding spatial precision of *azo*-HU308, we applied targeted illumination of three different square regions-of-interest on the plate (**Fig. 1G**). We observed [Ca^2+^]_i_ increases only in the cells that were irradiated, while cells that were outside the irradiation area did not respond (**Fig. 1H**). Combined, these results demonstrate that *azo*-HU308 is a photoswitchable ligand that robustly stimulates Ca^2+^ in INS-1 β-cells when activated with UV-light and isomerized to *cis*.

### *Azo*-HU308’s Effect on β-cell Ca^2+^ is Not Mediated by CB2R

After verifying the robust effect of *cis*-*azo*-HU308 to stimulate INS-1 cell [Ca^2+^]_i_, we wanted to determine its mechanism of action. Our previous study showed that *azo*-HU308 and HU308 elevate [Ca^2+^]_i_ in AtT-20(CB2) cells through CB2R and phospholipase C (PLC)-mediated Ca^2+^-release from intracellular stores^43^. Therefore, we co-applied CB2R antagonists alongside *azo*-HU308 to verify the contributions of CB2R signaling. To our surprise, *azo*-HU308 still caused a large Ca^2+^ response in INS-1 cells upon irradiation, even when co-applied with three different CB2R antagonists—AM630, SR144528, and JWH133 (**Fig. 2A and Fig. 2B**)^48–50^. Although HU308 is touted as a selective ligand for CB2R^42^, it is possible that our probe has off-target effects at other CBRs. INS-1 cells express multiple CBRs including CB1R and GPR55.^16,46^ However, co-application of the CB1R antagonist rimonabant^51^, or the GPR55 antagonist CID16020046^52^ failed to block *azo*-HU308’s photoswitching effect (**Fig. 2A, Fig. S3A, and Fig. S3B**). These results suggests that the effects of *azo*-HU308 in β-cells are not mediated by CB2R, or through off-target CBR pathways.

**FIGURE 2.**
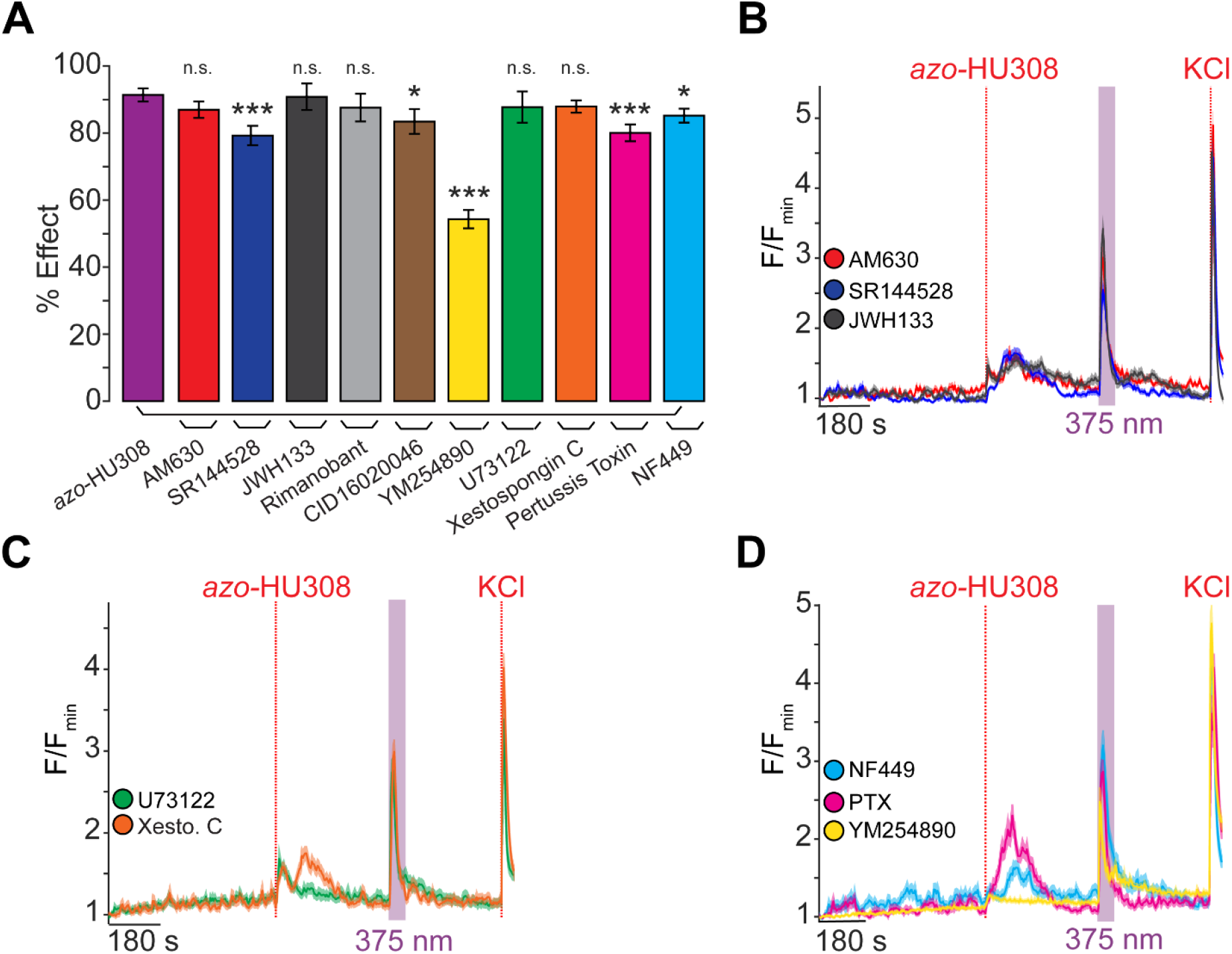
*Azo*-HU308’s photoswitching effect in INS-1 cells is not mediated by CB2R or G protein signaling. INS-1 cells were transfected with R-GECO and Ca^2+^ levels recorded by confocal microscopy. *Azo*-HU308 (20 μM) and 375 nm light were applied in conjunction with various signaling antagonists. Shown are: (**A**) Summary bar graph showing that the % Ca^2+^ effect for azo-HU308 photoswitching in the presence of CB2R inhibitors AM630 (20 μM; N = 151, T = 3), SR144528 (20 μM; N = 159, T = 3), and JWH133 (32 μM; N = 166, T = 3), which failed to block the *azo*-HU308’s effect. Similar results were observed with the CB1R inhibitor Rimanobant (2 μM; N = 103, T = 2), GPR55 inhibitor CID16020047 (20 μM; N = 104, T = 2), PLC inhibitor U73122 (10 μM; N = 89, T = 2), or the IP_3_R inhibitor Xestospongin C (1 μM; N = 152, T = 3). Inhibitors for G protein signaling, including G_s_-protein inhibitor NF449 (10 μM; N = 173, T = 3), G_i/o_-protein inhibitor pertussis toxin (100ng/mL; N = 135, T = 4), and G_q/11_ protein inhibitor YM524890 (10 μM; N = 120, T = 3) also failed to block *azo*-HU308 photoswitching (**B**) Averaged Ca^2+^ traces for CB2R inhibitors AM630 (red), SR144528 (blue), and JWH133 (black). **(C)**Average traces for U73122 (green) and Xestospongin C (orange). (**D**) Average Ca^2+^ traces for G protein inhibitors: G_s_-inhibitor NF449 (cyan), G_i/o_-inhibitor pertussis toxin (magenta) and G_q/11_-inhibitor YM254890 (yellow). Error bars = mean ± s.e.m. *P < 0.05, **P < 0.01, ***P < 0.001, ns = not significant.

Next, we asked whether PLC and Ca^2+^-release from internal stores were being stimulated by *cis-azo*-HU308, as we found in our previous study^43^. Surprisingly, both the PLC inhibitor U73122, or inositol 1,4,5-trisphosphate receptor (IP_3_R) inhibitor Xestospongin C failed to block the photoswitching effect (**Fig. 2A,C**), indicating that an entirely different Ca^2+^ signaling mechanism was involved.^53,54^ We tested several inhibitors of G protein signaling. The G_q/11_ G protein inhibitor YM254890 produced only a modest inhibition of the photoswitching effect, but the response was still largely intact (**Fig. 2A and Fig. 2D**, yellow).^55^ We applied the G_s_ protein inhibitor NF449 (**Fig. 2A and Fig. 2D**, cyan) and G_i/o_ protein blocker pertussis toxin (PTX, (**Fig. 2A and Fig. 2D**, magenta).^56,57^ Bodh drug caused only a small reduction of *cis*-*azo*-HU308’s effect on [Ca^2+^]_i_. Combined, these results suggest that the mechanism of action for *azo*-HU308’s effect on [Ca^2+^]_i_ release in INS-1 cells was independent of CB2R, other CBRs, or even GPCR activation entirely. These results suggest an entirely unique mechanism compared to our previous study in CB2R-overexpressing AtT-20 cells.

### *Azo-*HU308’s photoswitching effect is modulated by ion channel targets

Besides GPCRs, pancreatic β-cell [Ca^2+^]_i_ is affected by numerous transmembrane ion channels, including transient receptor potential (TRP) channels.^58,59^ Based on the rapid and transient nature of *azo*-HU308’s effect, we hypothesized that these may be involved as an unintended receptor target. To this end, we tested the effect of 2-aminoethoxydiphenyl borate (2-APB), which is a non-selective TRP channel inhibitor and IP_3_ receptor modulator,^60–62^ and observed a dose-dependent inhibition of *azo*-HU308 photoswitching (**Fig. 3A,B**). At low concentrations of 2-APB (1 μM, red) we observed a potentiation of the effect of *trans-azo*-HU308, as a larger [Ca^2+^]_i_ increase was observed upon probe addition, while the cis-effect remained constant. At a mid-range concentration (50 μM, blue) the photoswitching effect was significantly reduced, and at the highest concentration (100 μM, black), the effect of *trans*- and *cis*-*azo*-HU308 was completely abolished. We then conducted our Ca^2+^ imaging experiments in Ca^2+^-free extracellular solution (with added ethylene glycol-bis(β-aminoethyl ether)-N,N,N′,N′-tetraacetic acid (EGTA)) where we observed that *azo-*HU308’s photoswitching effect on Ca^2+^ was completely abolished (**Fig. 3A,C**). Combined, these results indicate that the Ca^2+^ response of *cis*-*azo*-HU308 is mediated by ion channel expressed on the INS-1 cell surface, and that the rise in [Ca^2+^]_i_ requires extracellular Ca^2+^ influx.

**FIGURE 3.**
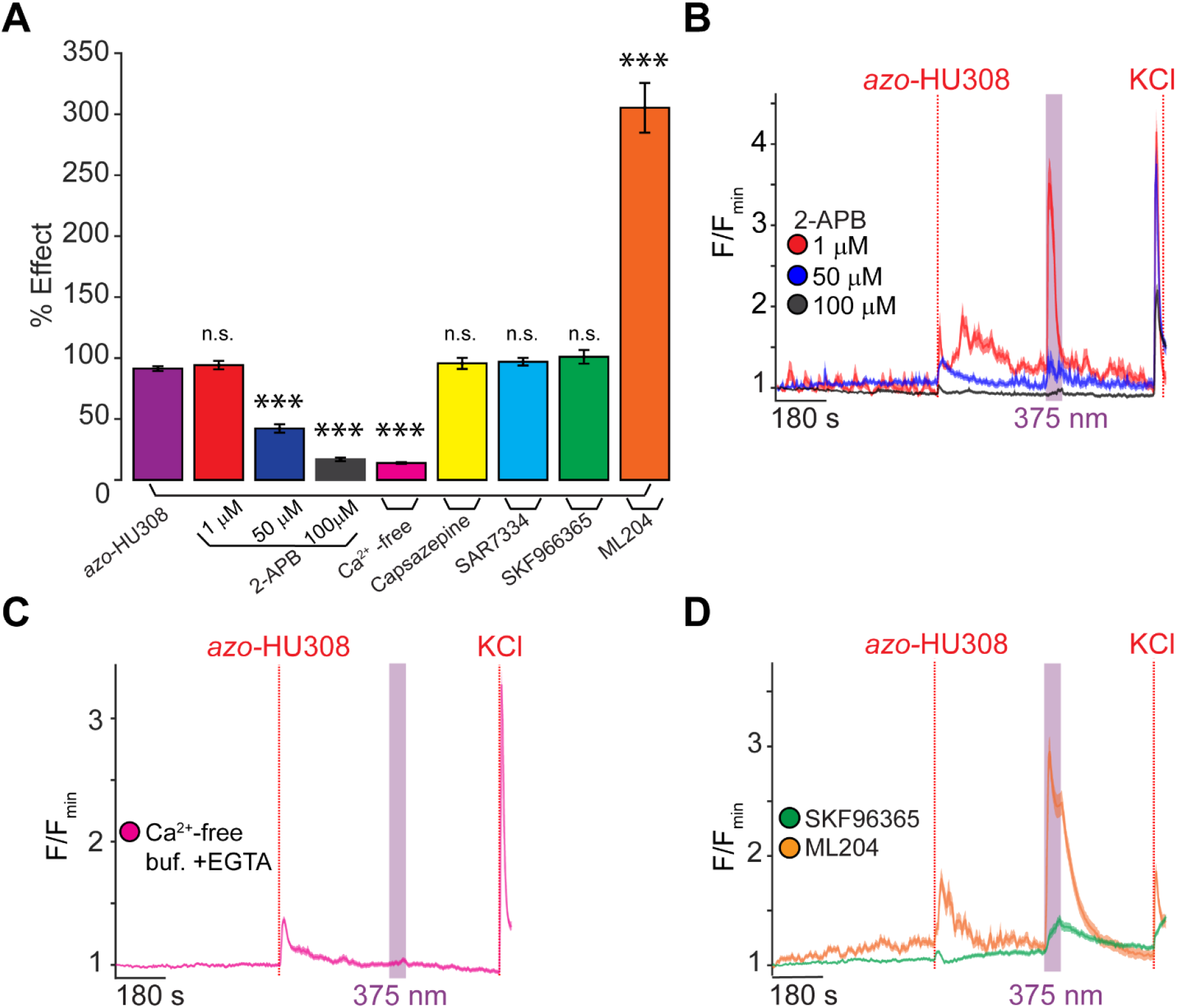
*Azo*-HU308’s effect on Ca^2+^ is modulated by pharmacological inhibitors of TRP channels. INS-1 cells were transfected with R-GECO and Ca^2+^ levels recorded by confocal microscopy. *Azo*-HU308 (20 μM) was applied in conjunction with various antagonists. (**A**) Summary bar graph showing a dose-responsive photoswitching effect on Ca^2+^ with 2-APB at different concentrations (1 μM; N = 100, T = 2), (50 μM; N = 93, T = 2), and (100 μM; N = 96, T = 2), and the response was completely abolished by the removal of extracellular Ca^2+^ using EGTA (0.1 mM) in the buffer (N = 200, T = 4). The TRPV1 blocker Capsazepine (5 μM; N = 119, T = 2), TRPC6 blocker SAR7334 (20 μM; N = 164, T = 3), broadband TRPC blocker SKF963665 (20 μM; N = 151, T = 3), and TRPC4 blocker ML204 (100 μM; N = 133, T = 3) failed to prevent the photoswitching effect. (**B**) Averaged Ca^2+^ traces showing dose-dependent inhibition of photoswitching response with 2-APB at 1 μM (red), 50 μM (blue), and 100 μM (black). (**C**) Averaged Ca^2+^ trace showing that the photoswitching effect requires an extracellular source of Ca^2+^ (magenta), as azo-HU308 photoswitching had no effect in the presence of EGTA. **(D)** Averaged Ca^2+^ traces showing the effect of SKF96365 (green); and that ML204 (orange) potentiated photoswitching. Error bars = mean ± s.e.m. *P < 0.05, **P < 0.01, ***P < 0.001, ns = not significant.

We then set out to determine the specific ion channels involved. INS-1 cells express a variety of TRP channels, which are known to respond to lipid agonists^58,63–65^. Transient receptor potential vanilloid 1 (TRPV1) is expressed in INS-1 cells^66^ and has been shown to be activated by lipids. However, when INS-1 cells were pretreated with the TRPV1 antagonist capsazepine^67^, *azo*-HU308 photoswitching effects remained intact (**Fig. 3A and Fig.S4A**), ruling out TRPV1’s role. Next, we set our sights on TRPC channels. Human and murine pancreatic islets and β-cells, including INS-1 cells, express multiple TRPC channels which are important for store-operated Ca^2+^ homeostasis and can regulate of insulin-secretion^59,68^. Gratifyingly, when we pre-treated with the broadband TRPC inhibitor SKF96365, there was a reduction in amplitude to the photoswitching effect of *azo*-HU308 (**Fig. 3A,D**, green)^69^. We then applied more selective antagonists. Interestingly, co-application of the TRPC4 inhibitor ML204 potentiated the photoswitching response (**Fig. 3A and Fig.3D**, orange). Additionally, the TRPC6 inhibitor SAR7334 failed to block the photoswitching response (**Fig. 3A and Fig. S4B**)^70,71^.

### TRPC channels mediate *azo-*HU308’s photoswitching effect on Ca^2+^ in INS-1 cells

Recent studies have shown that TRPC channels expression and regulation of insulin secretion from human and murine islets,^72,73^ and our broadband TRPC inhibitor SKF96365 reduced the amplitude of *azo*-HU308 photoswitching effects in INS-1 cells. Therefore, we focused on TRPC channels using blockers that have greater selectivity. Intriguingly, treatment with YM58483— which blocks TRPC3 and TRPC5 channels^74^—completely abolished the [Ca^2+^]_i_ response caused by *cis*-*azo*-HU308 (**Fig. 4A and Fig. 4B, red**). Next, we treated INS-1 cells with the selective TRPC3-specific inhibitor Pyr10^75^. In this case, we saw a ~50% reduction in the Ca^2+^ response caused by photoswitching (**Fig. 4A and Fig. 4B, blue**), demonstrating that TRPC3 may be involved but is not solely responsible for the effect. With both YM58483 and Pyr10 effectively reducing the *azo*-HU308 mediated Ca^2+^ response during irradiation, these results suggest that the proposed target of our photoswitch is likely through TRPC channels expressed in INS-1 cells.

**FIGURE 4.**
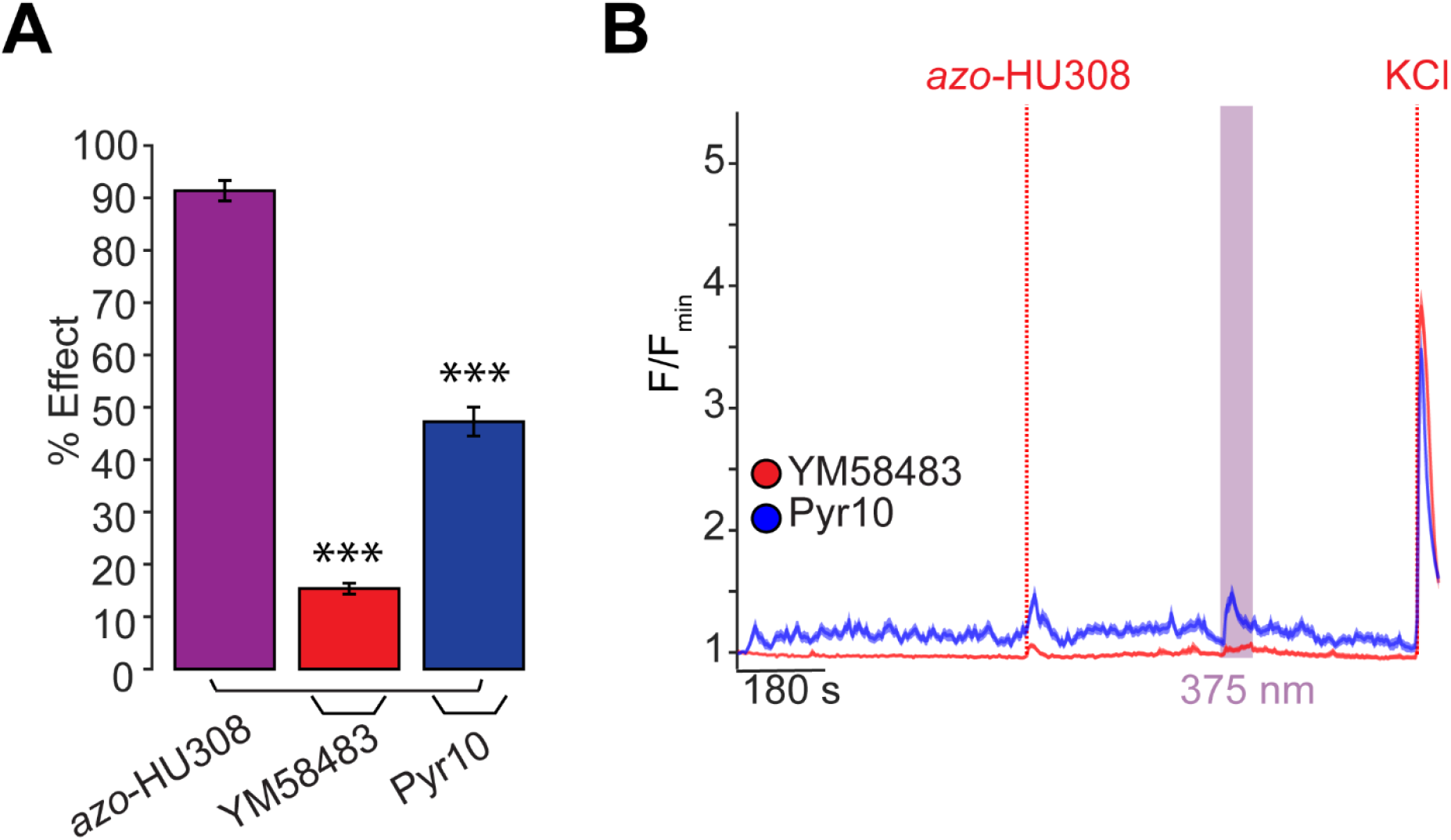
TRPC channels are a target for *azo*-HU308 in INS-1 cells. INS-1 cells were transfected with R-GECO and Ca^2+^ levels recorded by confocal microscopy. *Azo*-HU308 (20 μM) was applied in conjunction with TRPC channel antagonists. (**A**) Summary bar graph showing that TRPC channel inhibitors YM58483 (20 μM; N = 174, T = 4) and Pyr10 (15 μM; N = 200, T = 4) significantly reduced the photoswitching effect of *azo*-HU308. (**B**) Averaged Ca^2+^ trace showing that treatment with the TRPC3 and TRPC5 inhibitor YM58483 (red) eliminated the Ca^2+^ photoswitching response, while the TRPC3-selective antagonist Pyr10 (blue) reduced the Ca^2+^ photoswitching response by ~50%. Shaded error bars = mean ± s.e.m. *P < 0.05, **P < 0.01, ***P < 0.001, ns = not significant.

## DISCUSSION

This study demonstrates that the photoswitchable cannabinoid *azo*-HU308 robustly stimulates [Ca^2+^]_i_ in INS-1 cells when activated by UV-light, and that the probe is more active in the *cis-*configuration. While our first study using *azo*-HU308 in CB2R-overexpressing AtT20 cells showed that the probe triggers a prolonged Ca^2+^ response via Ca^2+^-induced Ca^2+^-release after CB2R and PLC activation^43^; here, an extensive pharmacological screen ruled-out this mechanism of action. In INS-1 cells, the intense and short-lived Ca^2+^ increase induced by *cis*-azo-HU308 is initiated solely by the influx of extracellular Ca^2+^, suggesting that an ion channel mechanism is involved. Our results also suggested that the plasma membrane ion channel target rapidly desensitizes, which led us to investigate the role of TRP ion channels. β-cells express a multitude of TRP channels permit the flux of Ca^2+^ ions into the cell, and we observed that the effect of *azo*-HU308 was modulated by a several TRP ion channels modulators, including classical non-selective TRP channel blockers like 2-APB and SKF96365. Gratifyingly, selective inhibitors for TRPC channels blocked azo-HU308’s effect, with the most significant inhibition coming from the TRPC3 and TRPC5 antagonist YM58483. While we have not entirely ruled out that azo-HU308 activates other targets in INS-1 cells, our results indicate that *azo*-HU308 is likely acting via TRPC channels. While these findings reveal an ion-channel mechanism for *azo*-HU308’s ability to robustly stimulate Ca^2+^ release in INS-1 cells, this study was performed in an *in vitro* system using immortalized β-cells. Future studies to investigate the action of *azo*-HU308 on human β-cells or cultured islets will be of interest.

In summary, we describe *azo*-HU308’s ability to optically stimulate Ca^2+^ levels with high spatiotemporal control through an ion channel mechanism. These findings shed light on non-GPCR pathways that can be activated by photoswitchable cannabinoids and sets the stage for future applications of photoswitchable cannabinoids in β-cells to reveal how cannabinoid drugs regulate β-cell excitability and Ca^2+^ homeostasis.

## ABBREVIATIONS

ATP: Adenosine triphosphate
CB2R: Cannabinoid 2 receptor
GPCR: G protein-coupled receptor
GLP1R: Glucagon-like peptide-1 receptor
GSIS: Glucose-stimulated insulin secretion
IP_3_R: Inositol triphosphate receptor
K_ATP_: ATP-sensitive K^+^-channel
N: Number of cells
PLC: Phospholipase
C ROI’s: Regions of interest
T: Trials, number of biological replicates
T2DM: Type II diabetes mellitus

## ACKNOWLEDGEMENTS

The authors thank Erick Carreira, Miroslav Kosar, and Roman Sarott for synthesizing *azo*-HU308 compound used in these experiments. We also thank Carsten Schultz for providing microscopy resources.

