## Supporting Information - Figures and Tables for "The Photoswitchable Cannabinoid *Azo*-HU308 Enables Optical Control of Ca^2+^ Signaling in Pancreatic β-Cells via a Non-CB2 TRPC Channel Mechanism"

### SUPPORTING FIGURES

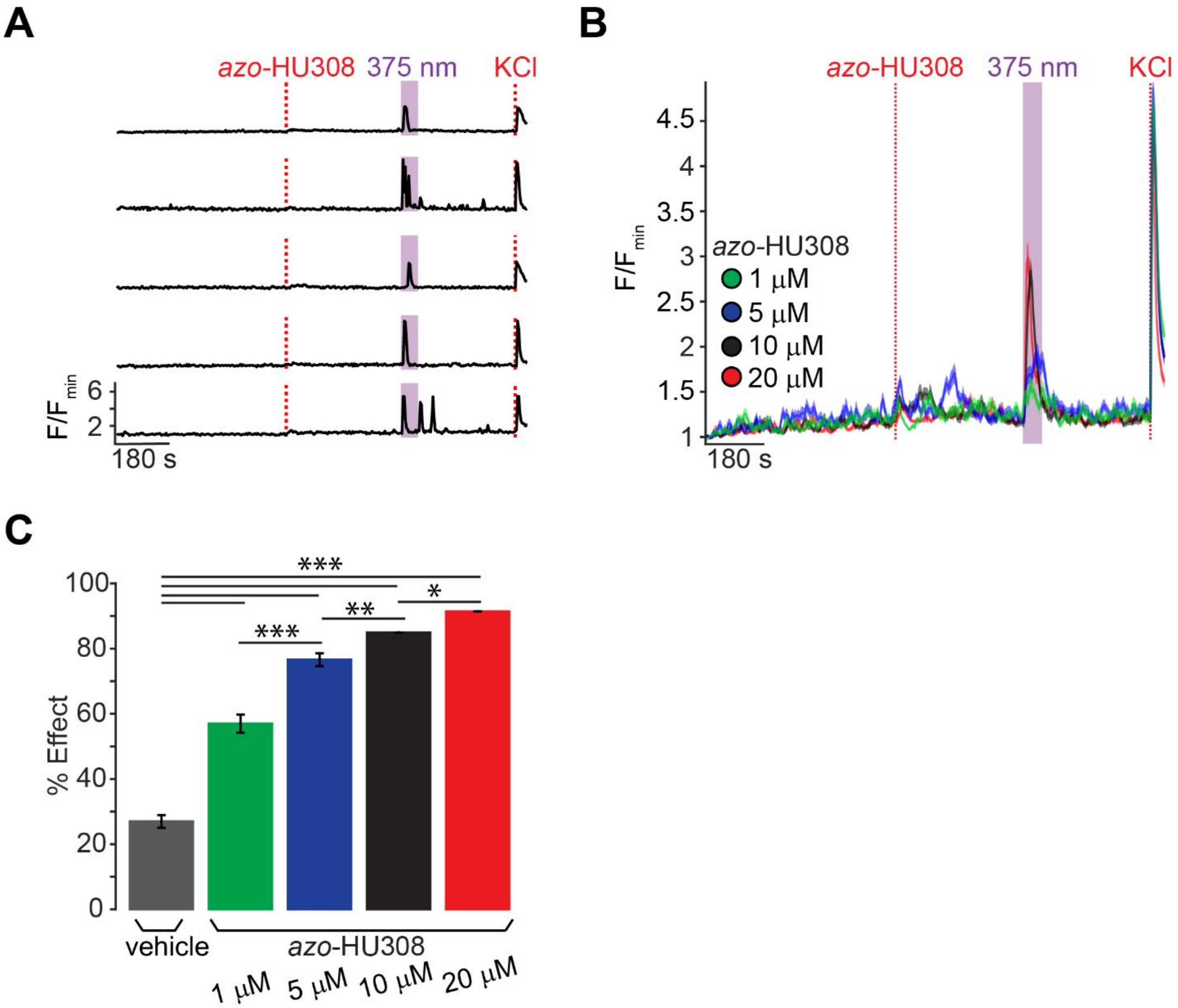


**FIGURE S1. *azo*-HU308 increases $Ca^{2+}$ transient in INS-1 cells with stimulated by UV-A light.** INS-1 cells were transfected with RGECO and their $Ca^{2+}$ levels monitored by confocal microscopy. (**A**) Representative single cell $Ca^{2+}$ traces for INS-1 cells after the addition of *azo*-HU308 (20 μM), which caused an increase in $Ca^{2+}$ after photoswitching to *cis* with 375 nm light. KCl (30 mM) was added at the end of each experiment to maximize intracellular $Ca^{2+}$.
(**B**) Averaged $Ca^{2+}$ imaging traces showing dose-response for *azo*-HU308 at 1μM (N = 157, T = 3), 5 μM (N = 162 cells, T = 3), 10 μM (N = 149, T = 3), and 20 μM (N = 170, T = 4) followed by photoactivation with 375 nm light. (**C**) Summary bar graph comparing Ca2+ responses to the vehicle (N = 203, T = 4) and dose-response of *azo*-HU308 as shown in part B. Error bars = mean ± s.e.m. *P < 0.05, **P < 0.01, ***P < 0.001, ns = not significant = P > 0.05.

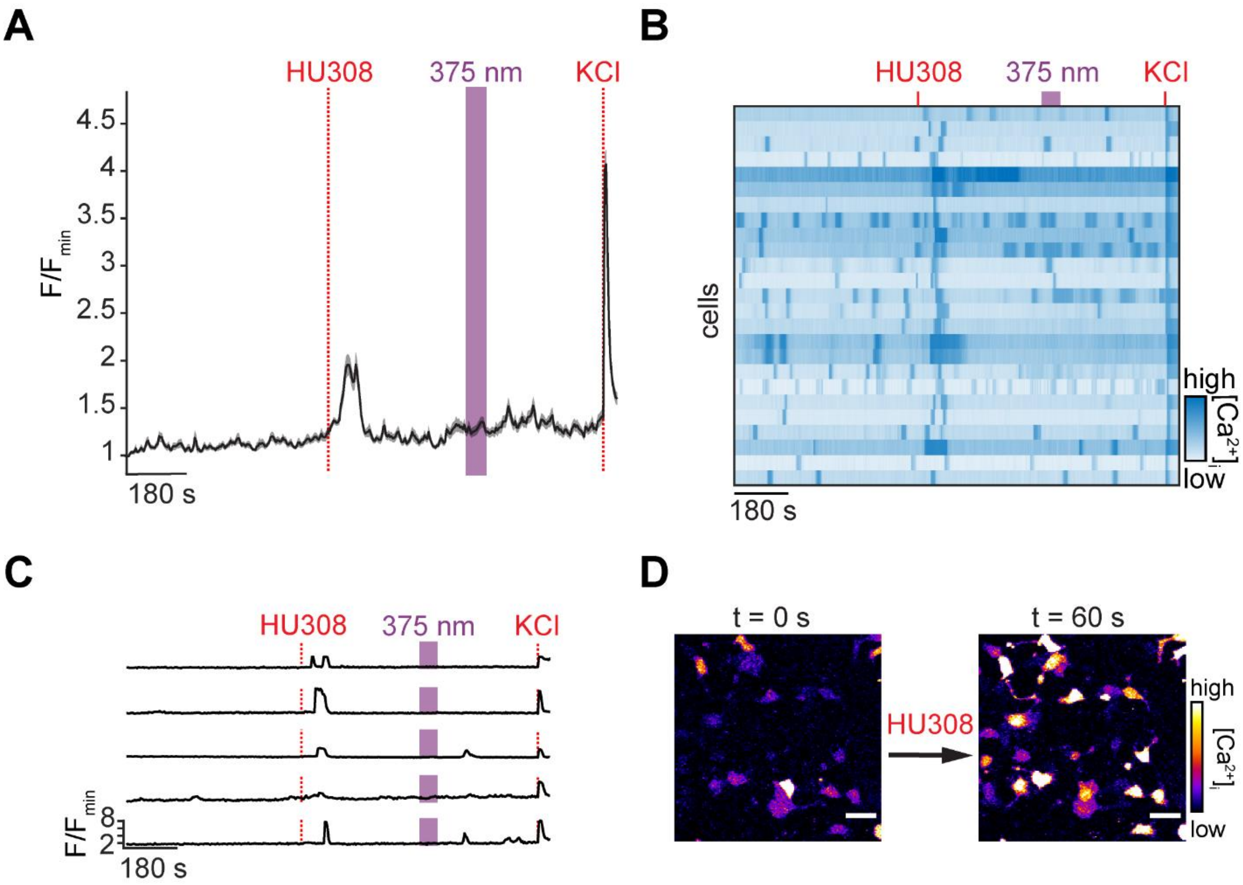


**FIGURE S2. Non-photoswitchable HU308 addition causes a short $Ca^{2+}$ transient in INS-1 cells.** INS-1 cells were transfected with RGECO and their $Ca^{2+}$ levels monitored by confocal microscopy. (**A**) Averaged (N = 150, T = 3), (**B**) heatmap of 25 cells, and (**C**) representative single cell $Ca^{2+}$ traces for INS-1 cells after the addition of HU308 (20 μM). An increase in $Ca^{2+}$ was observed upon addition, but no response was triggered upon 375 nm irradiation. For A, shaded error bars = mean ± s.e.m.. For B, $Ca^{2+}$ traces from 25 representative cells were normalized to the KCl response. (**D**) Representative fluorescence micrographs showing increased $Ca^{2+}$ in INS-1 cells after HU308 addition. Scale bar = 30 μm.

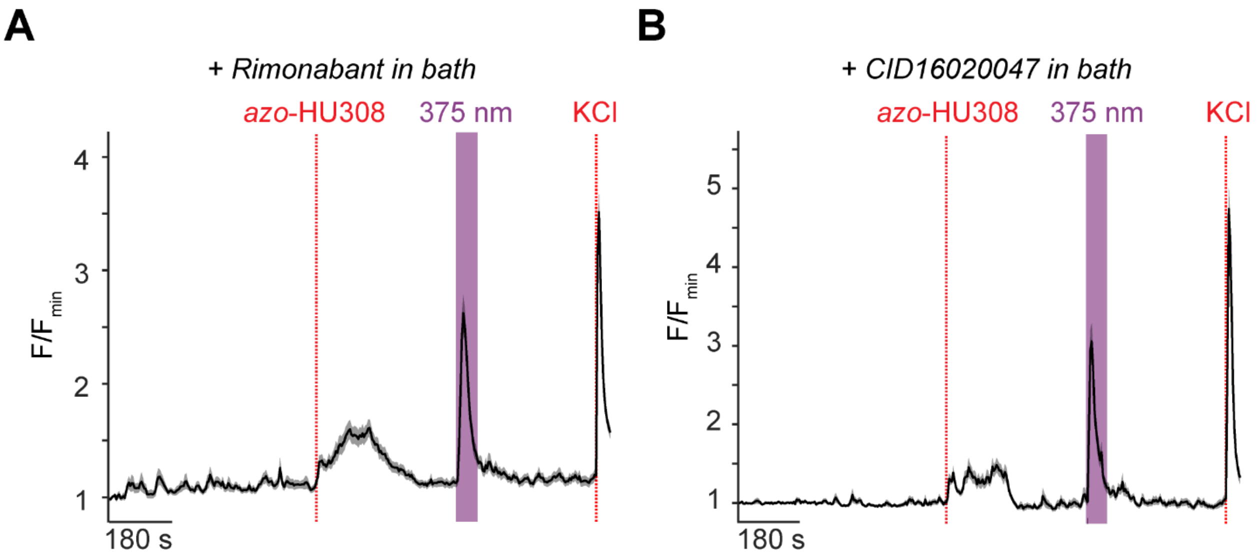


**FIGURE S3. *azo*-HU308's photoswitching effect on $Ca^{2+}$ is not CB1- or GPR55-mediated.** INS-1 cells were transfected with RGECO and their $Ca^{2+}$ levels monitored by confocal microscopy. (**A,B**) Averaged $Ca^{2+}$ imaging traces showing the effect of *azo*-HU308 (20 μM) addition and 375 nm photostimulation in the presence of (A) the CB1 antagonist rimonabant (2 μM, N = 103, T = 2) and (B) the GPR55 antagonist CID16020047 (20 μM, N = 104, T = 2). A $Ca^{2+}$ increase was still observed on addition and stimulation. Shaded error bars = mean ± s.e.m.

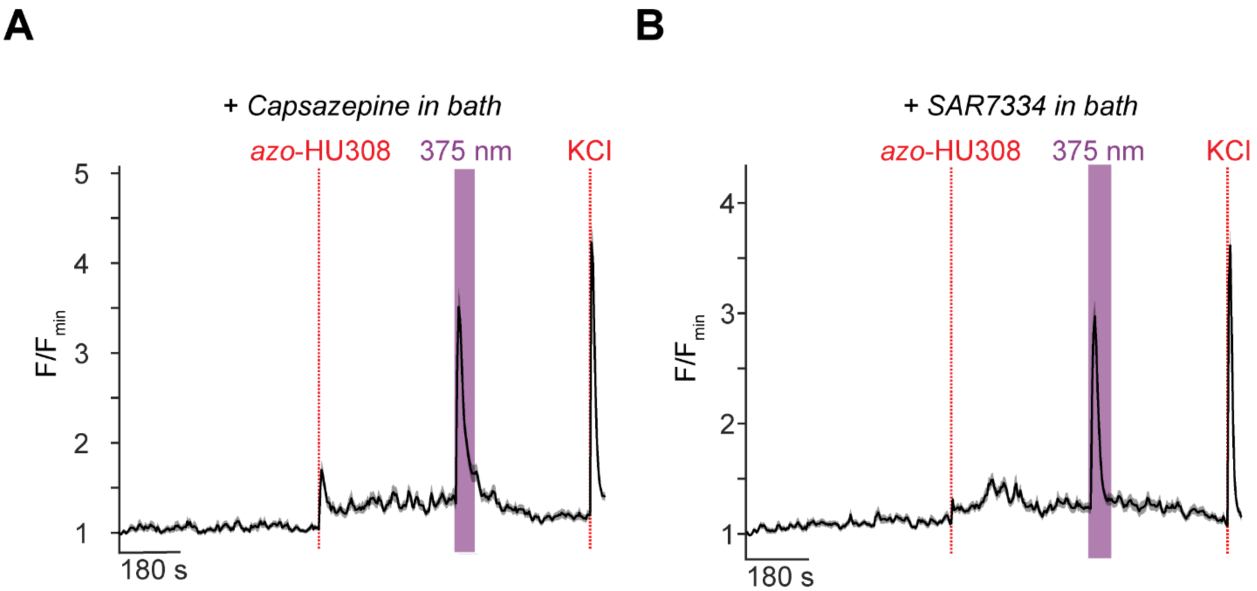


**FIGURE S4. *azo*-HU308's photoswitching effect is not TRPV1- or TRPC6-mediated.** INS-1 cells were transfected with RGECO and their $Ca^{2+}$ levels monitored by confocal microscopy. (**A,B**) Average $Ca^{2+}$ imaging traces showing that *azo*-HU308 addition and photostimulation with 375 nm light still had an effect on $Ca^{2+}$ in the presence of (A) the TRPV1 blocker Capsazepine (5 μM, N = 119, T = 2) and (B) the TRPC6 blocker SAR7334 (20 μM, N = 164, T = 3). Shaded error bars = mean ± s.e.m.

### SUPPORTING TABLES

#### TABLE S1: Statistical significance calculations for Figure 1 bar graph

| conditions | p-value | significance |
|---|---|---|
| *trans-azo*-HU308 / *cis-azo*-HU308 | 5.32E-68 | *** |
| HU308 + 375 nm / *cis-azo*-HU308 | 1.55E-48 | *** |
| vehicle + 375 nm / *cis azo*-HU308 | 8.41E-74 | *** |
| vehicle + 375 nm / HU308 + 375 nm | 0.0238 | * |
| HU308 dark / HU308 + 375 nm | 2.47E-07 | *** |
| HU308 dark / *cis-azo*-HU308 | 1.80E-11 | *** |
| vehicle dark / *cis-azo*-HU308 | 0.1169 | n.s. |
| vehicle dark / HU308 + 375 nm | 5.91E-11 | *** |
| vehicle dark / vehicle + 375 nm | 0.0139 | * |

***P<0.005 (dark blue, *P<0.05 (light blue), ns = P>0.05 (white)

#### TABLE S2: Statistical significance calculations for Figure 2 bar graph

| conditions | p-value | significance |
|---|---|---|
| *cis-azo*-HU308 / *cis azo*-HU308 + AM630 | 0.1564 | n.s. |
| *cis-azo*-HU308 / *cis azo*-HU308 + CID16020047 | 0.0378 | * |
| *cis-azo*-HU308 / *cis azo*-HU308 + YM254890 | 6.62E-25 | *** |
| *cis-azo*-HU308 / *cis azo*-HU308 + JWH133 | 0.8940 | n.s. |
| *cis-azo*-HU308 / *cis azo*-HU308 + NF449 | 0.0305 | * |
| *cis-azo*-HU308 / *cis azo*-HU308 + Pertussis Toxin | 3.71E-04 | *** |
| *cis-azo*-HU308 / *cis azo*-HU308 + Rimonabant | 0.3640 | n.s. |
| *cis-azo*-HU308 / *cis azo*-HU308 + SR144528 | 4.83E-04 | *** |
| *cis-azo*-HU308 / *cis azo*-HU308 + U73122 | 0.4006 | n.s. |
| *cis-azo*-HU308 / *cis azo*-HU308 + Xestospongin C | 0.1982 | n.s. |

***P<0.01 (dark blue, *P<0.05 (light blue), ns = P>0.05 (white)

#### TABLE S3: Statistical significance calculations for Figure 3 bar graph

| conditions | p-value | significance |
|---|---|---|
| *cis-azo*-HU308 / *cis azo*-HU308 + 100μM 2-APB | 1.74E-75 | *** |
| *cis-azo*-HU308 / *cis azo*-HU308 + 1μM 2-APB | 0.4480 | n.s. |
| *cis-azo*-HU308 / *cis azo*-HU308 + 50μM 2-APB | 3.03E-31 | *** |
| *cis-azo*-HU308 / *azo*-HU308 + $Ca^{2+}$ free | 1.06E-133 | *** |
| *cis-azo*-HU308 / *cis azo*-HU308 + Capsazepine | 0.3447 | n.s. |
| *cis-azo*-HU308 / *cis azo*-HU308 + ML204 | 1.80E-26 | *** |
| *cis-azo*-HU308 / *cis azo*-HU308 + SAR7334 | 0.1265 | n.s. |
| *cis-azo*-HU308 / *cis azo*-HU308 + SKF966365 | 0.0829 | n.s. |

***P<0.01 (dark blue, *P<0.05 (light blue), ns = P>0.05 (white)

**TABLE S4: Statistical significance calculations for Figure 4 bar graph**

| conditions | p-value | significance |
|---|---|---|
| *cis-azo*-HU308 / *cis azo*-HU308 + Pyr10 | 1.26E-30 | *** |
| *cis-azo*-HU308 / *cis azo*-HU308 + YM58483 | 1.58E-113 | *** |

***P<0.01 (dark blue, *P<0.05 (light blue), ns = P>0.05 (white)

**TABLE S5: Statistical significance calculations for SI Figure 1 bar graph**

| conditions | p-value | significance |
|---|---|---|
| *cis* 10μM *azo*-HU308 / *cis* 20μM *azo*-HU308 | 0.0227 | n.s. |
| vehicle + 375 nm / *cis* 10μM *azo*-HU308 | 3.36E-61 | *** |
| *cis* 1μM *azo*-HU308 / *cis* 5μM *azo*-HU308 | 2.01E-08 | *** |
| vehicle + 375 nm / *cis* 1μM *azo*-HU308 | 8.02E-18 | *** |
| vehicle + 375 nm / *cis* 20μM *azo*-HU308 | 8.41E-74 | *** |
| *cis* 5μM *azo*-HU308 / *cis* 10μM *azo*-HU308 | 0.0032 | *** |
| vehicle + 375 nm / *cis* 5μM *azo*-HU308 | 1.48E-50 | *** |

***P<0.01 (dark blue, *P<0.05 (light blue), ns = P>0.05 (white)
